# Muscle progenitor cells are required for the regenerative response and prevention of adipogenesis after limb ischemia

**DOI:** 10.1101/2020.02.24.963777

**Authors:** Hasan Abbas, Lindsey A. Olivere, Michael E. Padgett, Cameron A. Schmidt, Brian F. Gilmore, Kevin W. Southerland, Joseph M. McClung, Christopher D. Kontos

## Abstract

Peripheral artery disease (PAD) is nearly as common as coronary artery disease, but few effective treatments exist, and it is associated with significant morbidity and mortality. Although PAD studies have focused on the vascular response to ischemia, skeletal muscle cells play a critically important role in determining the phenotypic manifestation of PAD. Here, we demonstrate that genetic ablation of Pax7^+^ muscle progenitor cells (MPCs, or satellite cells) in a murine model of hind limb ischemia (HLI) resulted in a complete absence of normal muscle regeneration following ischemic injury, despite a lack of morphological or physiological changes in resting muscle. Compared to ischemic muscle of control mice (Pax7^WT^), the ischemic limb of Pax7-deficient mice (Pax7^Δ^) was unable to generate significant force 7- or 28-days after HLI in *ex vivo* force measurement studies. A dramatic increase in adipose infiltration was observed 28 days after HLI in Pax7^Δ^ mice, which replaced functional muscle. To investigate the mechanism of this adipogenic change, mice with inhibition of fibro/adipogenic precursors (FAPs), another pool of MPCs, were subjected to HLI. Inhibition of FAPs decreased muscle adipose fat but increased fibrosis. MPCs cultured from mouse muscle tissue failed to form myotubes *in vitro* following depletion of satellite cells *in vivo*, and they displayed an increased propensity to differentiate into fat in adipogenic medium. Importantly, this phenotype was recapitulated in patients with critical limb ischemia (CLI), the most severe form of PAD. Skeletal muscle samples from CLI patients demonstrated an increase in adipose deposition in more ischemic regions of muscle, which corresponded with a decrease in the number of satellite cells in those regions. Collectively, these data demonstrate that Pax7^+^ MPCs are required for normal muscle regeneration after ischemic injury, and they suggest that targeting muscle regeneration may be an important therapeutic approach to prevent muscle degeneration in PAD.

## Introduction

Peripheral artery disease (PAD) caused by atherosclerosis of the peripheral arteries affects over 200 million individuals globally and is a major contributor to disease burden in both developing and developed countries (1, 2). Current treatment options are limited to surgical and percutaneous revascularization approaches (3), both of which have minimal impact on long-term morbidity and mortality (4, 5). The clinical course of PAD ranges from the milder manifestation of intermittent claudication (IC), resulting in pain with ambulation that resolves with rest, to the more severe critical limb ischemia (CLI), characterized by pain at rest with or without tissue necrosis (3). Although CLI affects only 10-15% of patients with PAD, it is responsible for a substantial health care burden, as these patients often progress to limb amputation and have significantly greater morbidity and mortality (6, 7). Although therapeutic approaches to PAD all target revascularization and tissue perfusion, it has been observed that patients with similar degrees of atherosclerotic vascular occlusion often present with markedly different severity of disease (8, 9).

Recent evidence from our group and others supports the idea that skeletal muscle responses to tissue ischemia, and not solely the vascular supply, play an important role in determining the muscle response to limb ischemia (9–12). In mice subjected to hind limb ischemia (HLI), a model of PAD, the genetic background strongly influences outcomes. For example, C57BL/6 mice display not only robust angiogenesis but also a muscle regenerative response that typically leads to full recovery from HLI. In stark contrast, HLI in BALB/c mice typically results in muscle degeneration and auto-amputation, and even muscle that does survive fails to recover function, i.e., force generation (9). Although this genetic difference was previously attributed to differences in collateral vessel density (13–16), muscle progenitor cells (MPCs) isolated from these strains of mice display markedly different responses to experimental ischemia *in vitro*, independent of blood supply, which is consistent with a muscle cell-specific determinant of the response to ischemia and reflects the differential responses observed *in vivo*. While these findings demonstrate that tissue perfusion is not the only factor that determines severity of ischemic muscle injury, the mechanisms by which skeletal muscle responds to ischemia remain unclear. A genetic variant in at least one gene, *Bag3*, has been linked to this differential ischemic response in mice (12), but it is not known whether these effects are at the level of mature muscle cells or MPCs.

MPCs, commonly known as satellite cells, lie between the basal lamina and plasma membrane of skeletal muscle cells and are critical regulators of postnatal myofiber regeneration (17, 18). MPCs are defined by expression of the Pax3 homolog Pax7, and they are the primary cells that serve as a unipotent stem cell population for myogenesis following injury (19). However, these cells have a limited capacity for self-renewal, and repeated replication cycles may result in depletion of the satellite cell pool (20). The development of genetically modified mouse models to ablate MPCs has allowed investigation of the role of Pax7^+^ MPCs in various disease states. In particular, mice inducibly expressing *Diphtheria* toxin A (DTA) only in Pax7^+^ cells have been used to demonstrate a requirement for these MPCs in muscle regeneration in a variety of conditions. Most of these studies have been performed using cytotoxic injury models, such as cardiotoxin, freeze injury, or BaCl_2_ injury (21). However, it is known that different modes of injury have unique characteristics. For example, glycerol injury results in a more adipogenic phenotype compared to other modes of injury (22). Although we and others have begun to characterize the skeletal muscle response to ischemia (9, 10, 12, 23), the role of MPCs in this process remains unexplored.

In addition to MPCs, the discovery of a novel subpopulation of fibro/adipogenic progenitors (FAPs) in mature skeletal muscle (24) has led to considerable focus on the role of these cells in pathological skeletal muscle conditions. FAPs isolated from skeletal muscle were able to cause white fat infiltration in diseased but not healthy muscle because myofibers have a significant inhibitory effect on the differentiation of FAPs (25). This suggested a contribution from the environment in determining the fate of these cells. FAP expansion has also been shown to regulate the MPC pool during muscle regeneration in addition to playing a critical role in skeletal muscle homeostasis (26).

Here, we used genetically modified mouse models to explore the role of Pax7^+^ MPCs in the skeletal muscle response to hindlimb ischemia. We demonstrate a near complete absence of skeletal muscle regeneration after HLI in mice following ablation of Pax7^+^ satellite cells. Furthermore, ischemic, Pax7-deficient muscle displayed a dramatic increase in adipogenesis that was driven at least in part by FAPs. Consistent with these findings in mice, decreased MPC numbers and increased adipogenesis were observed in more ischemic regions of skeletal muscle of CLI patients. These findings demonstrate the requirement for Pax7^+^ MPCs in ischemic skeletal muscle regeneration, and they provide important new insights into the pathogenesis of PAD.

## Results

To study the role of Pax7^+^ MPCs in a mouse model of PAD, we crossed Pax7-Cre^ERT2^ mice to ROSA26^DTA^ mice. To ablate satellite cells, we injected tamoxifen (Pax7^Δ^) or corn oil as a control (Pax7^WT^) for 5 days, followed by HLI surgery. To maintain MPC ablation, mice were fed a diet supplemented with either corn oil or tamoxifen. To validate the model, muscle sections from the non-ischemic tibialis anterior (TA) muscle were stained for the satellite cell marker Pax7, and in the tamoxifen-treated mice there was a complete absence of satellite cells (Fig. S1), demonstrating successful ablation of all satellite cells within skeletal muscle.

### Satellite cell ablation does not alter resting muscle morphology or physiology

To examine whether satellite cell ablation resulted in changes in resting muscle, we isolated the TA muscle from the contralateral, non-ischemic limbs of Pax7^Δ^ and Pax7^WT^ mice after HLI and compared the skeletal muscle histologically by H&E staining. Muscle morphology and architecture looked identical in Pax7^Δ^ and Pax7^WT^ mice (Fig. 1A). Furthermore, *ex vivo* force generation of extensor digitorum longus (EDL) muscle did not differ between Pax7^Δ^ and Pax7^WT^ mice (Fig. 1B), demonstrating that absence of satellite cells does not alter resting muscle physiology. Lastly, we examined whether deletion of the endogenous skeletal muscle progenitor cell pool affects muscle fiber type distribution. Staining of both type 1, slow-twitch fibers, which are highly oxidative, and glycolytic type IIa and IIb fibers (which are more abundant in TA muscle) demonstrated no differences between groups of mice, demonstrating that loss of Pax7^+^ MPCs does not cause a shift in myofiber metabolism at rest (Fig. 1C).

**Figure 1.**
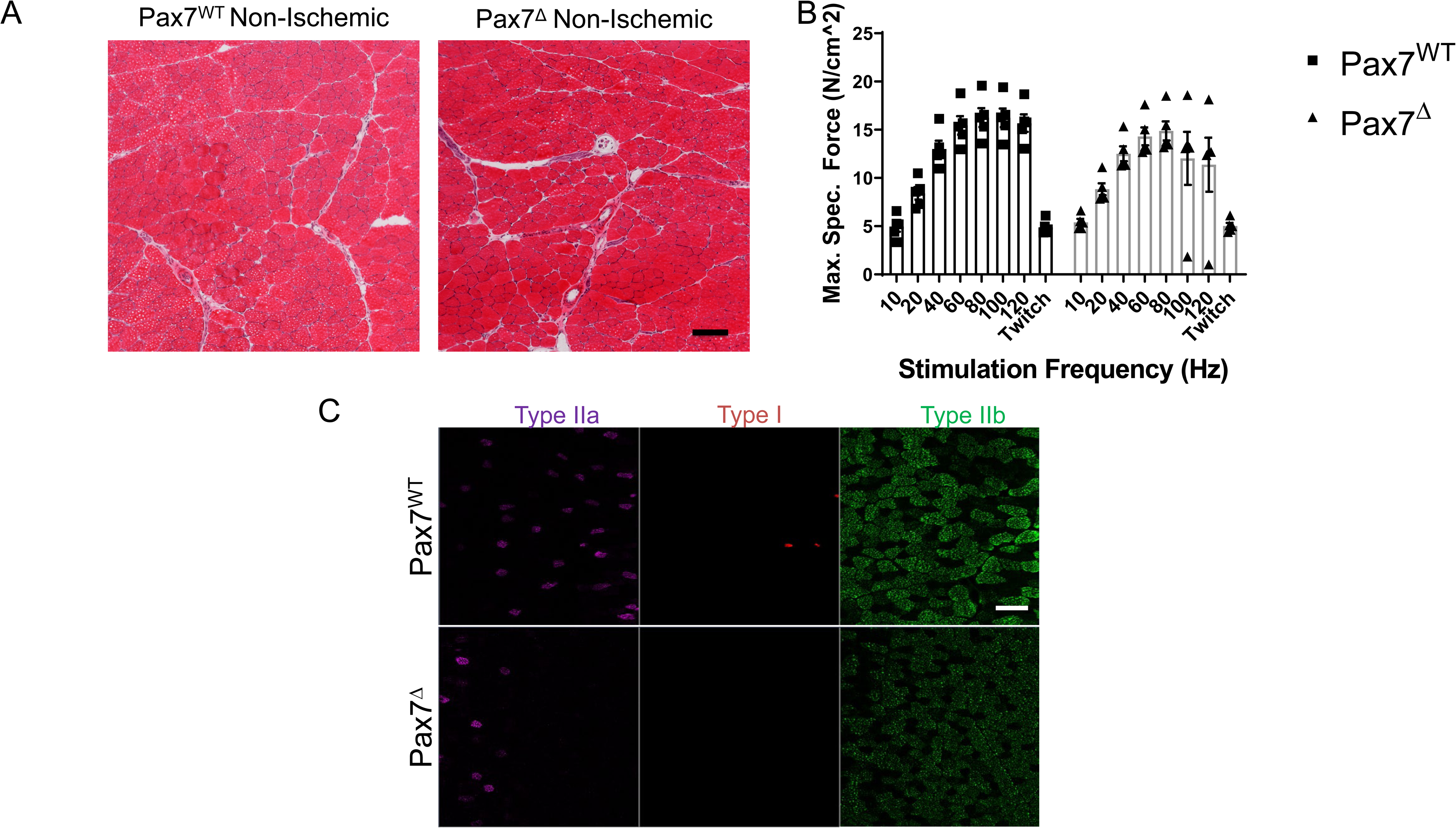
Pax7^+^ MPC ablation does not alter resting muscle morphology one week after ischemia. (A) H&E stains of skeletal muscle from mice with ablated satellite cells are not distinguishable from muscle with intact satellite cells (n=3-4 per group). (B) EDL muscles from mice (n=5 per group) were isolated, and their ability to generate force was measured on a force transducer. Ablation of satellite cells did not impair the ability of resting skeletal muscle to generate force. (C) Pax7^+^ MPC ablation did not alter resting muscle fiber type distribution one week after ischemia (n=3-4 per group). Scale bar=100 μm.

### Satellite cell ablation in ischemic muscle results in complete absence of regeneration one week after ischemia

To determine the effect of MPC ablation after ischemia, Pax7^Δ^ and Pax7^WT^ mice were subjected to unilateral HLI and examined 7 days later. Following ablation of satellite cells, markers of skeletal muscle regeneration (embryonic myosin heavy chain expression and centralized myonuclei) were absent in the ischemic limb of Pax7^Δ^ mice (Fig. 2A-C). To exclude the possibility that genetic ablation of MPCs with DTA had a non-specific effect on the vasculature, muscle sections were stained for the endothelial cell marker PECAM (CD31). Not only was the endothelium intact, but the total endothelial area relative to the muscle area was in fact increased in Pax7^Δ^ mice (Fig. 2D), suggesting a possible vascular compensation for the muscle loss. Satellite cell activation and proliferation normally occur after muscle injury in general and are observed after limb ischemia as well. One week after HLI, Pax7^WT^ mice displayed a significant increase in the number of Pax7^+^ cells in ischemic TA muscle in contrast to the contralateral, non-ischemic limb, confirming normal satellite cell activation in this model. As expected, this activation was absent in Pax7^Δ^ mice lacking satellite cells, consistent with their inability to regenerate muscle following ischemic injury (Fig. 2E, F).

**Figure 2.**
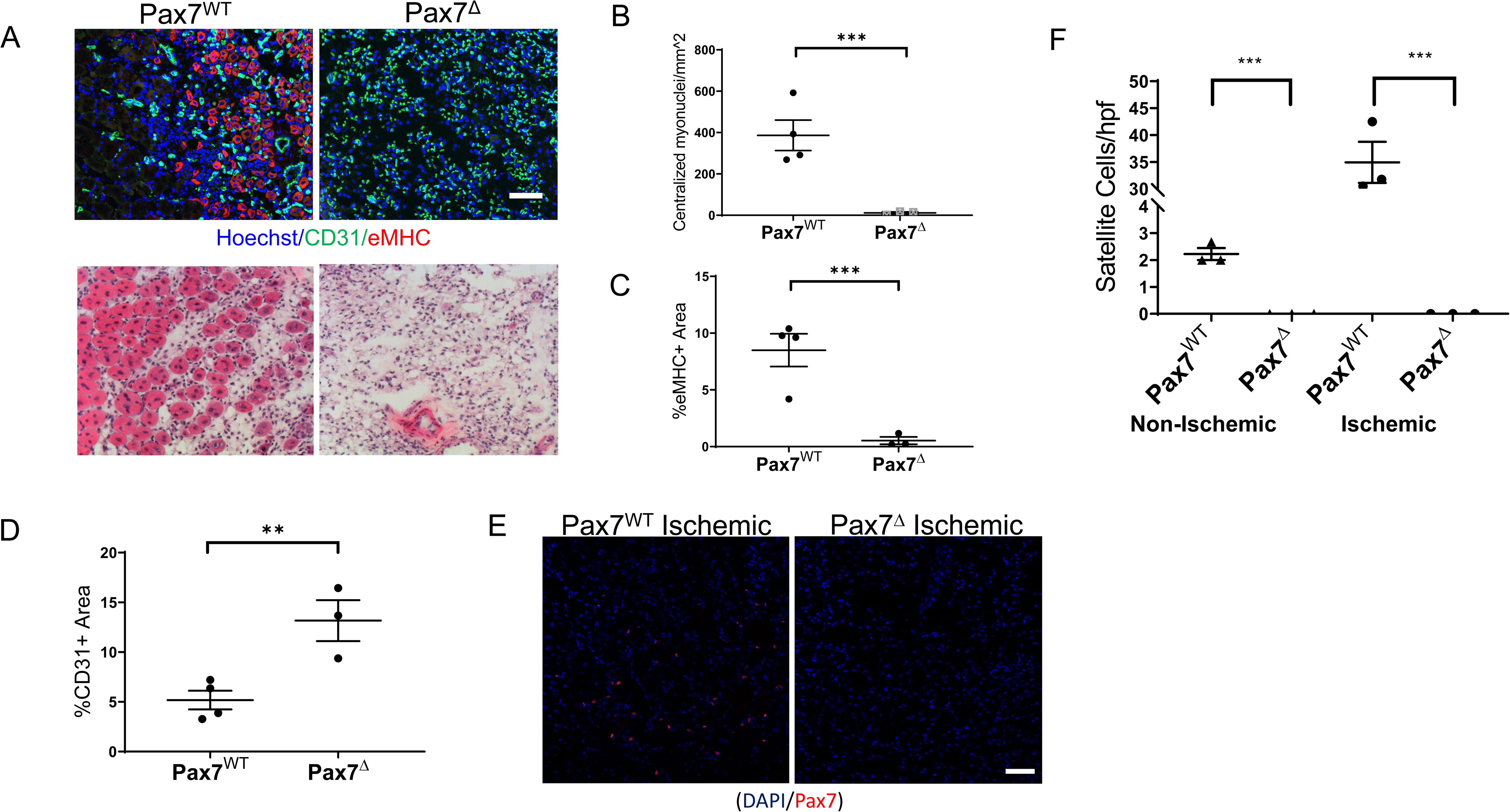
Ablation of Pax7^+^ MPCs in mice results in a complete lack of a muscle regenerative response one week after HLI. (A) Muscle regeneration was examined by staining for embryonic myosin heavy chain (eMHC, red) and endothelial cells (CD31, green), *top*, and for centralized myonuclei by H&E, *bottom*. (B-D) Quantification of centralized myonuclei (B) and eMHC (C) demonstrates a complete lack of regenerative response to ischemia. Quantification of CD31 area (D) demonstrates an increase in endothelial area relative to muscle area. (E, F) One week after HLI surgery, there was a significant increase in the number of Pax7^+^ cells per high power field in the ischemic TA of Pax7^WT^ mice but not in muscle of Pax7^Δ^ mice. Compared to resting muscle, there was a 10-15-fold increase in the number of Pax7^+^ cells in injured Pax7^WT^ muscle, consistent with activation of satellite cells following injury. Scale bar=100μm

### Chronic satellite cell ablation in ischemic muscle results in complete absence of regeneration one month after ischemia

To investigate the effects of satellite cell ablation on long-term muscle recovery from ischemia, Pax7^Δ^ and Pax7^WT^ mice were subjected to HLI and followed for 14 and 30 days after surgery. Consistent with responses observed in parental C57BL/6 mice, ischemic Pax7^WT^ mice displayed improved muscle architecture at day 14 post-HLI. Expression of eMHC had resolved by this time point, although there were still centralized myonuclei, and an inflammatory infiltrate was still present in the interstitial spaces between muscle fibers at this time point (Fig. 3A). These features were further improved by day 30, with near compete resolution of inflammation (Fig. 3B). In contrast, Pax7^Δ^ mice displayed a persistent absence of muscle regeneration with an accompanying increase in cellularity characteristic of a ongoing inflammation (Fig. 3A, B). Strikingly, muscle of late stage ischemic Pax7^Δ^ mice displayed a dramatic increase in adipose observed both histologically and by microCT (Fig. 3A, B, Fig. S2), which was also evidenced grossly by the inability of whole muscle tissue to sink in aqueous solution (Fig. S2A). Whereas distinct, individual myofibers were visualized by microCT in control muscle (Fig. S2B), EDL muscle from Pax7^Δ^ mice was markedly atrophied and displayed significant soft tissue adipogenic changes (Fig. S2B). These findings suggested that the chronic absence of satellite cells after ischemic injury resulted not only in a loss of muscle regeneration but also a shift in the cellular makeup of injured muscle. Persistent satellite cell ablation in Pax7^Δ^ mice 30 days after HLI was verified by Pax7 immuostaining (Fig. 3C, D). In the non-ischemic limb, satellite cell numbers were similar to the day 7 timepoint, whereas satellite cell number diminished significantly in the ischemic limb by day 30 (∼4/hpf compared to ∼35/hpf on day 7 post-HLI) and were only slightly higher than in the non-ischemic limb at this stage (Fig. 3C, D).

**Figure 3.**
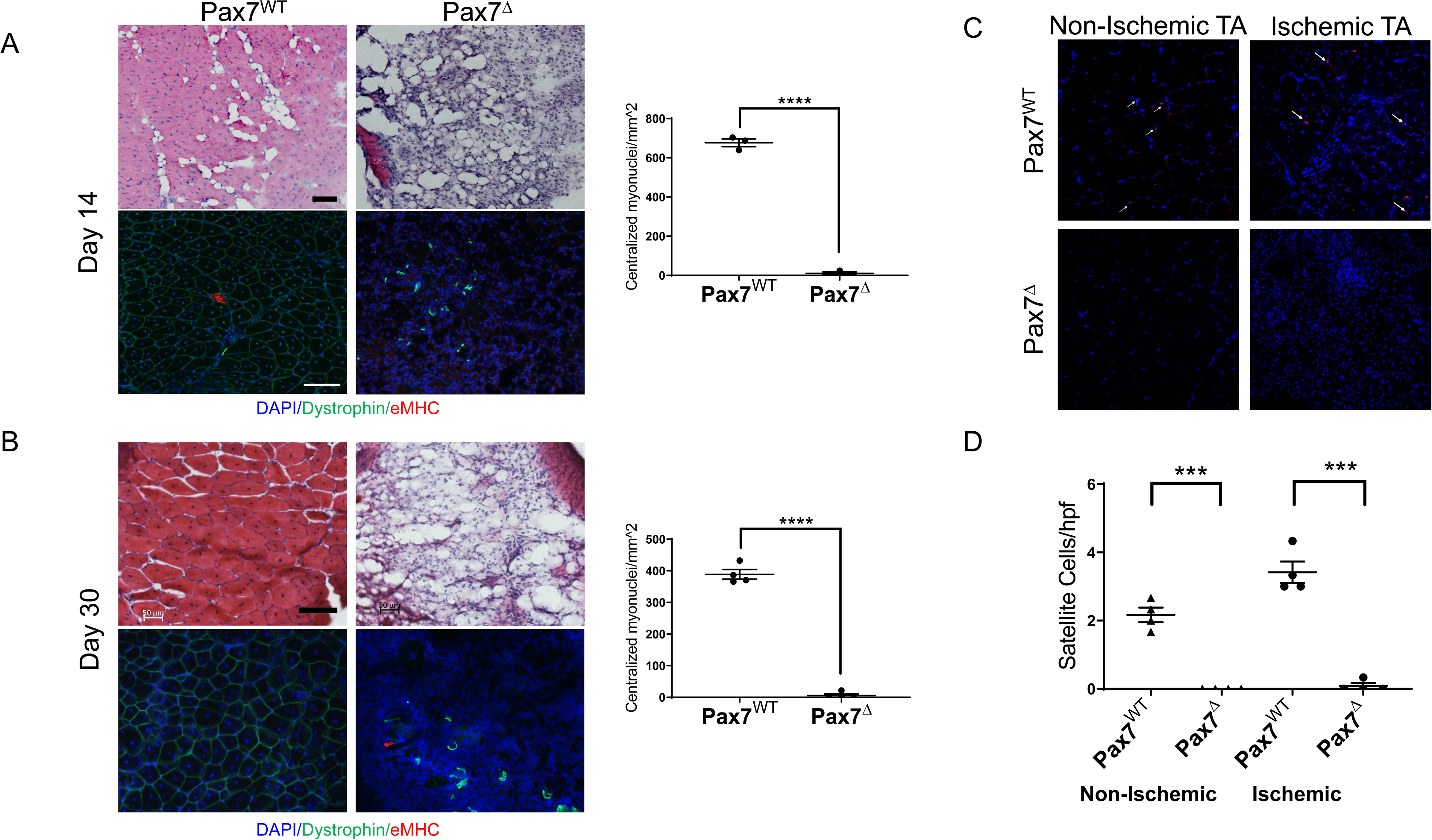
Sustained deletion of satellite cells results in long-term prevention of muscle regeneration after HLI. (A, B) H&E staining and quantification of regenerating fibers, demonstrated by myofibers with centralized myonuclei, of the ischemic TA muscle at 14 days (A) and 30 days (B) after ischemia demonstrated a complete lack of regeneration. (C, D) 30 days after HLI Pax7^+^ cells per high power field were significantly increased (2-fold) in the ischemic relative to the non-ischemic TA muscle of Pax7^WT^ mice, although their numbers were diminished compared to 7 days post-HLI. Pax7^+^ cells were persistently absent in Pax7^Δ^ mice. Scale bar=100 μm

### Long-term satellite cell ablation in ischemic muscle results in impaired force generation

Because long-term satellite cell ablation resulted in markedly abnormal muscle tissue morphology, we tested *ex vivo* muscle force generation to determine the functional effects of this injury. Force generation in Pax7^Δ^ and Pax7^WT^ mice correlated with histological findings, as there was a significant impairment in both maximal force generation and the time-tension force integral in EDL muscle of Pax7^Δ^ mice compared to that of Pax7^WT^ mice 3 days after ischemia (Fig. 4A, B). In stark contrast, force generation in the non-ischemic EDL mirrored that observed on day 7 post-HLI (Fig. 4C, D), confirming that resting skeletal muscle is unaffected by satellite cell ablation even after 30 days.

**Figure 4.**
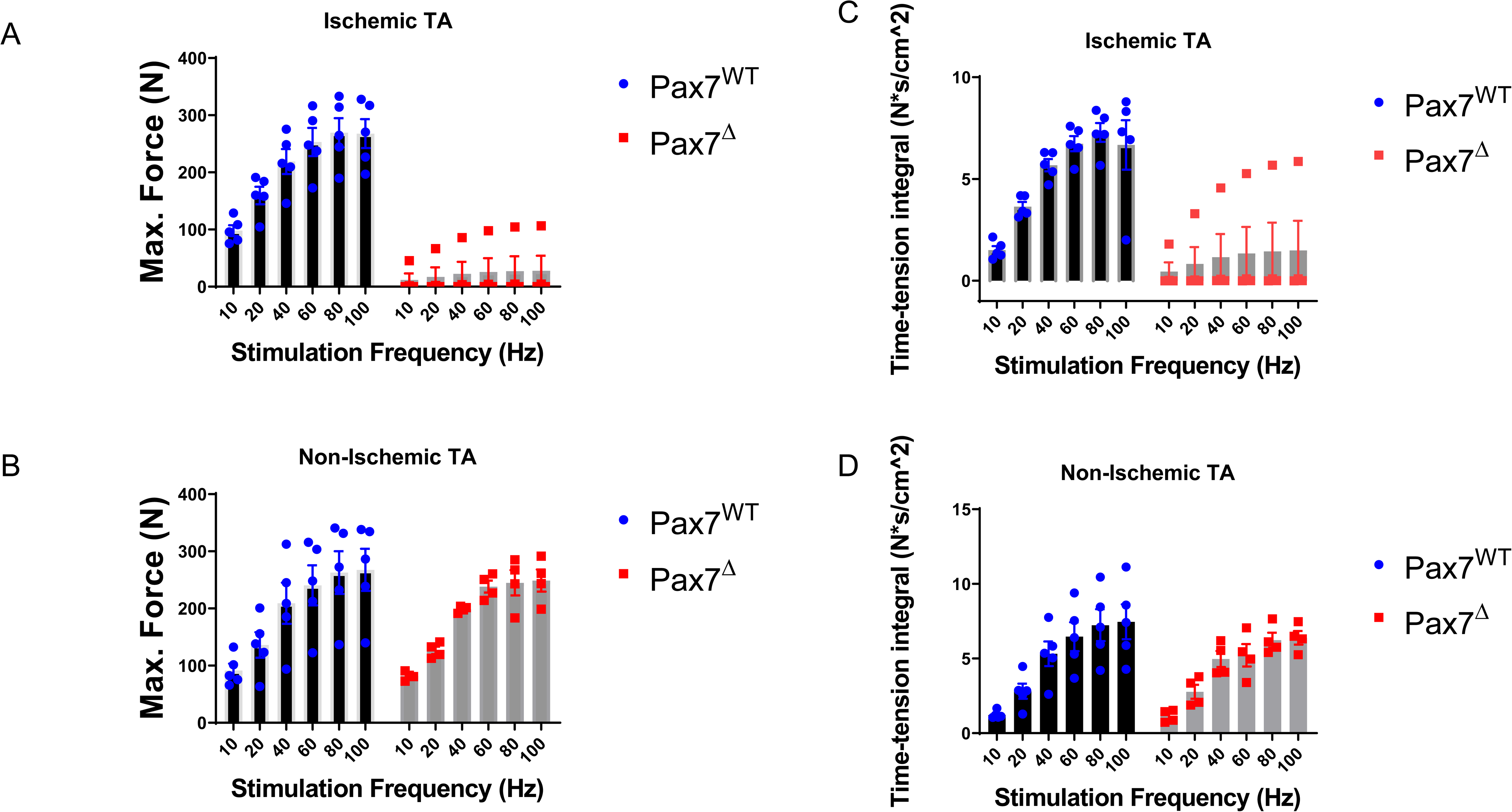
Pax7^+^ MPC ablation impairs force generation 30 days after HLI. (A) The maximum force generated by the ischemic EDL muscle was significantly lower in Pax7^Δ^ mice. (B) Maximum force was unchanged in the non-ischemic limb of Pax7^Δ^ mice. (C, D) The time-tension integral, a measure of work done in a single contraction, of muscle 3-days after HLI mirrored the maximum force data in both ischemic (C) and non-ischemic TA muscle (D) (n=4-5 per group).

### Ablation of Pax7^+^ MPCs in mice results in marked fat infiltration of skeletal muscle following ischemia

A key feature of muscle injury is that different modes of injury can result in varying regenerative responses. For example, unlike cardiotoxin-mediated injury, glycerol injection induces a more adipogenic change to the muscle (22). In contrast, the mdx mouse, a genetic model of muscular dystrophy, fails to accurately recapitulate many of the adipogenic changes observed in patients with muscular dystrophy. The lipid deposition seen in Pax7^Δ^ mice 30 days after HLI is reminiscent not only of that of patients with muscular dystrophy but also of patients with CLI (27). To investigate the adipogenic changes that occur in skeletal muscle following ischemic injury, we used two different complementary lipid stains, BODIPY 493/503 and oil red O, to examine fatty changes 7 days after HLI. Oil red O staining showed a small amount of fat deposition in the control Pax7^WT^ TA muscle, which was significantly increased in Pax7^Δ^ muscle, and these findings were mirrored by the BODIPY staining and quantification of the BODIPY stain (Fig. 5A). The increased fat deposition in Pax7^Δ^ muscle after long-term injury resulted in the need to cut thicker tissue (∼30 μm) sections, which also resulted in what appeared to be increased non-specific Oil Red O and BODIPY staining (data not shown). To overcome this issue, we immunostained for perilipin, which is selectively localized to the periphery of lipid droplets and thus specifically marks adipose accumulation. Perilipin staining also revealed a significant increase in adipogenesis in Pax7^Δ^ TA muscle compared to that of Pax7^WT^ mice 14 days post-HLI (Fig. 5B), and this difference was further enhanced by day 30 post-HLI (Fig. 5C). These findings suggest that any pathological adipogenesis is normally cleared during muscle regeneration after ischemia, but the lack of Pax7^+^ MPCs results in aberrant lipid accumulation, which may contribute to the pathogenesis of PAD.

**Figure 5.**
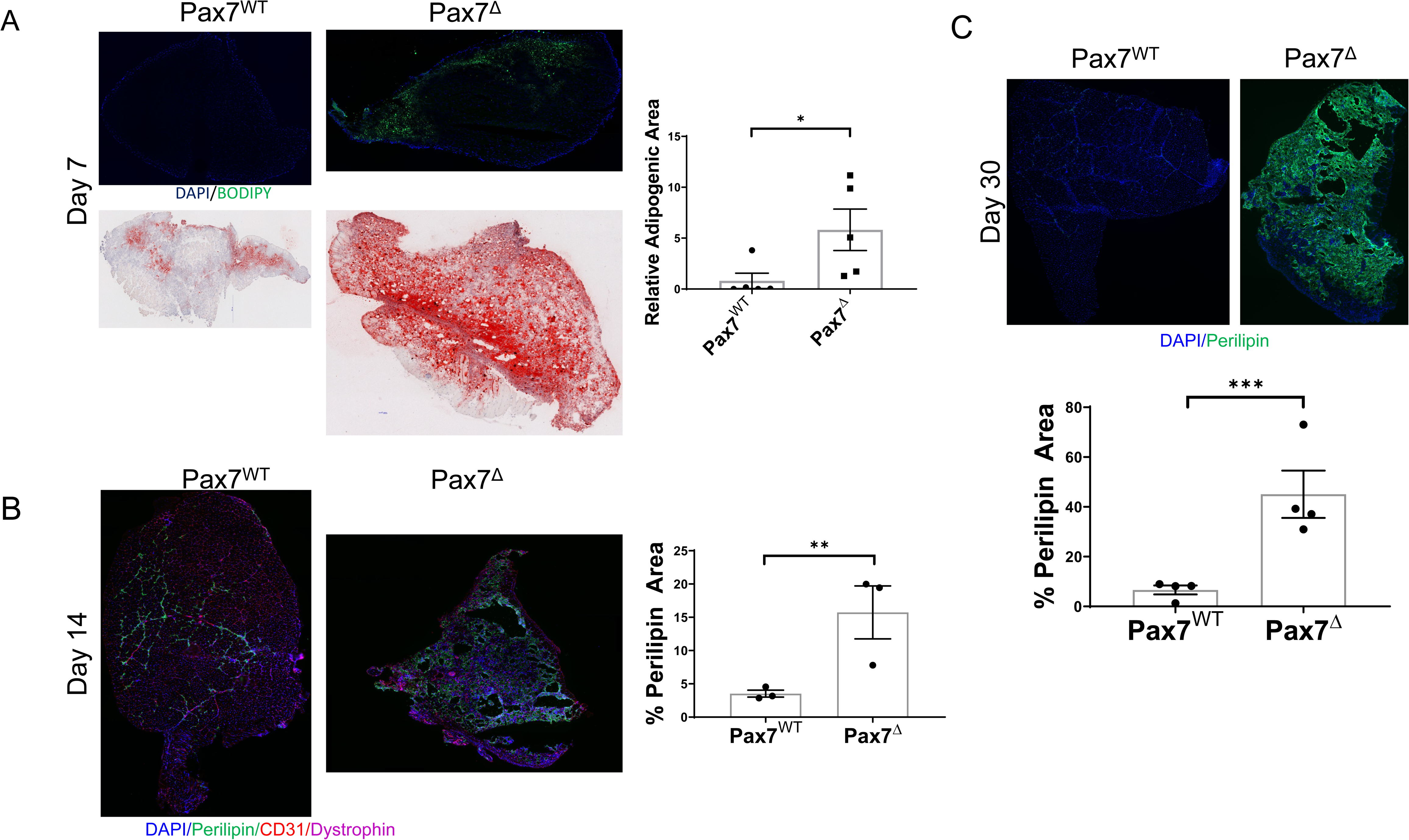
Ablation of Pax7^+^ MPCs in mice results in marked fat infiltration within skeletal muscle following ischemia. (A) BODIPY (*top*) and oil red O staining (*bottom*) of the ischemic TA muscle demonstrated significantly increased lipid staining in Pax7^Δ^ mice compared to Pax7^WT^ 7 days after HLI surgery. Quantification was performed on BODIPY-stained tissues. (B) Perilipin staining and quantification of adipose in the ischemic TA muscle demonstrated increased lipid staining in Pax7^Δ^ mice compared to Pax7^WT^ 14 days after HLI surgery. (C) Perilipin staining of the ischemic TA muscle 30 days after HLI surgery demonstrated increased lipid staining in Pax7^Δ^ mice compared to Pax7^WT^. Compared to day 14, the relative adipose area increased in Pax7^Δ^ muscle whereas it appeared to decrease slightly in control muscle.

### Batimastat, an MMP inhibitor that inhibits FAPs and adipogenesis, promotes fibrosis after ischemia in the absence of satellite cells

To begin to elucidate the origins of the adipogenic changes observed after ischemia in Pax7^Δ^ mice, we explored the potential contribution of fibro/adipogenic progenitor cells (FAPs) to the phenotype. FAPs have been shown to induce adipogenic changes in skeletal muscle in limb girdle muscular dystrophy type II (27) and in other pathological conditions (22). Batimastat is a non-specific MMP inhibitor that has been shown to inhibit adipogenesis in skeletal muscle by inhibiting FAPs (22, 27). We reasoned that if FAPs contribute to adipogenesis in HLI, then treating mice with batimastat should alleviate the degree of adipogenesis after HLI in the absence of satellite cells. Indeed, treatment of Pax7^Δ^ mice with batimastat during recovery from HLI resulted in significantly decreased adipogenic area compared to that observed in vehicle-treated Pax7^Δ^ mice (Fig. 6A). Notably, this change was accompanied by a corresponding increase in fibrosis (Fig. 6B). Collectively, these findings suggest that in the absence of satellite cells, ischemia drives FAPs to promote adipogenesis, which may play an important role in the pathophysiology of PAD.

**Figure 6.**
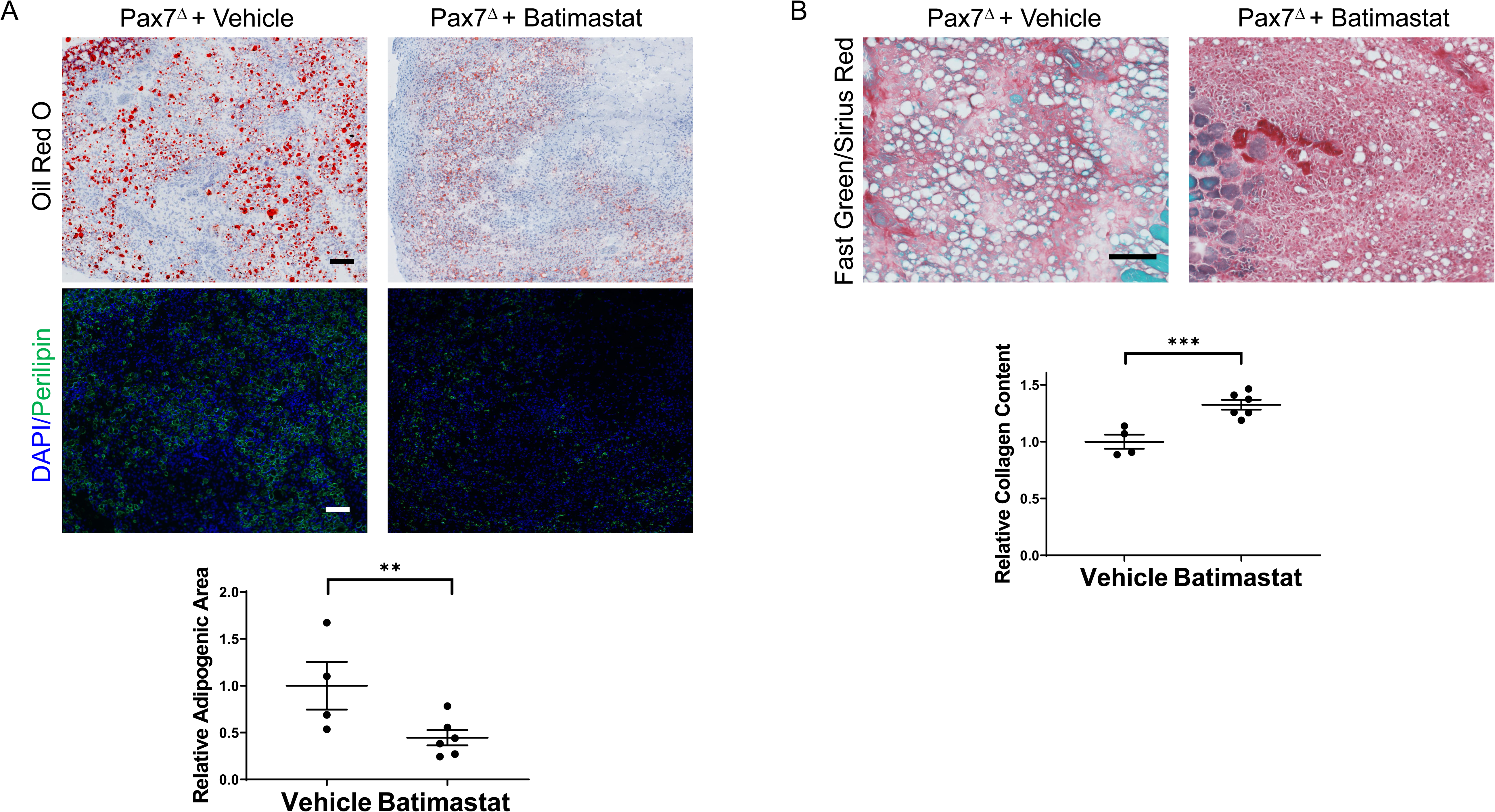
Inhibiting FAPs with batimastat reduces adipogenesis and increases fibrosis after HLI in the absence of satellite cells. (A) Batimastat treatment significantly decreased total fat in Pax7^Δ^ ischemic TA muscle as determined by oil red O and perilipin staining. (B) Fast Green/Sirius red staining demonstrated a corresponding significant increase in collagen content in Pax7^Δ^ ischemic TA muscle, consistent with a switch from adipogenesis to fibrosis after inhibition of FAPs (n=4-6 per group). Scale bars= 100 μm

### Isolation and differentiation of myoblasts following satellite cell ablation *in vivo* results in defective myogenesis and increased adipogenesis *in vitro*

It is well known that myoblasts isolated from whole muscle tissue retain their ability to differentiate and fuse into mature skeletal myotubes *in vitro*. Pax7^+^ satellite cells comprise a small percentage (<10%) of the mononuclear cells isolated from muscle tissue that have the potential to differentiate into muscle (i.e., MPCs). Once MPCs are isolated and plated *in vitro*, satellite cells rapidly lose expression of Pax7 and differentiate into MyoD-expressing committed myoblasts (28). Prior studies have demonstrated that deletion of Pax7 satellite cells *in vitro*, after plating, does not impair myoblast differentiation (29). To our knowledge, however, no studies have examined the effect of *in vivo* ablation of Pax7^+^ cells on subsequent myoblast differentiation *in vitro* and whether this might affect the propensity of cells to differentiate toward an adipogenic lineage. To test this, mice were treated with either tamoxifen or corn oil for 5 days to ablate Pax7^+^ cells *in vivo*, then muscle was harvested and mononuclear cells/myoblasts were isolated and plated *in vitro*. When cultured in muscle differentiation medium, only cells from Pax7^WT^ mice were able to form mature myotubes, as evidence by expression of the myogenic regulatory factor myogenin and myosin heavy chain (MHC) (Fig. 7A). To determine whether MPCs isolated from Pax7^WT^ or Pax7^Δ^ mice have an increased propensity to differentiate into adipocytes, cells were plated in adipogenic medium. Because increased adipogenesis in Pax7^Δ^ mice was observed *in vivo* in the setting of ischemia, cells were incubated for 12 days under hypoxic conditions to simulate ischemia. Compared to cells from Pax7^WT^ mice, cells isolated from Pax7^Δ^ mice had an increased propensity to form adipocytes, as demonstrated by oil red O staining (Fig. 7B). These findings suggest that in the absence of Pax7^+^ cells, Pax7^−^ cells with the potential to fuse and differentiate into muscle are driven toward an adipocyte lineage, although it is unclear whether these cells are FAPs or if they are derived some other progenitor cell population.

**Figure 7:**
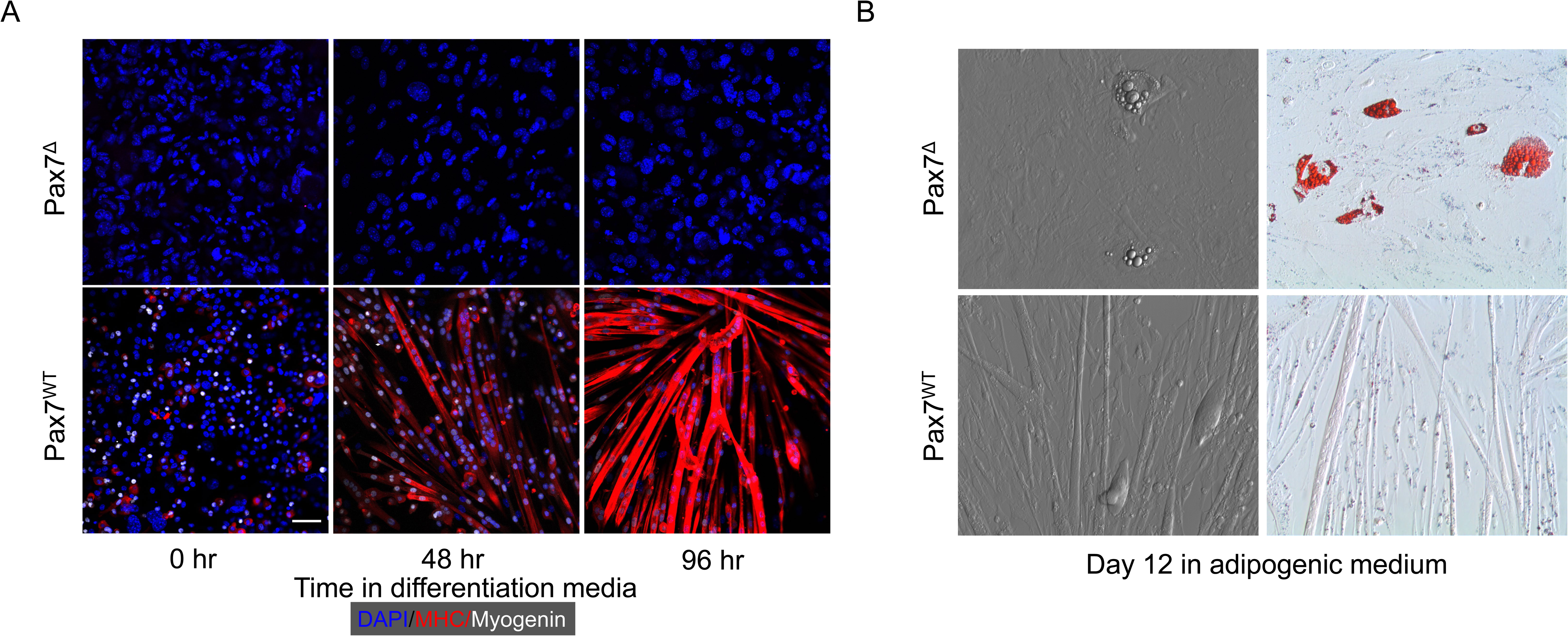
Myoblasts isolated from Pax7-depleted muscle fail to differentiate under hypoxic conditions and display increased adipogenesis. (A) Pax7^WT^ myoblasts in differentiation medium expressed myosin heavy chain (MHC) under hypoxia whereas Pax7^Δ^ cells failed to fuse and did not express the early differentiation marker myogenin or MHC. (B) When grown in adipogenic medium, Pax7^Δ^ myoblasts had a higher propensity to form oil red O^+^ lipid droplets. Similar results were observed in three independent experiments.

### Critical limb ischemia patients have increased adipogenesis and fewer satellite cells in regions of greater ischemia

To examine whether the adipogenic changes observed in our preclinical model are also seen clinically, we obtained skeletal muscle tissue from CLI patients undergoing limb amputation. In this setting, tissue that is farthest from the amputation site (distal) is typically the most ischemic, whereas proximal tissue, closer to the amputation site is less ischemic and often relatively healthy. Paired proximal and distal gastrocnemius muscle samples were obtained from 10 subjects, and adipose area was analyzed by perilipin staining. Distal, more ischemic muscle displayed significantly greater adipose area (Fig. 8A). Because the increase in adipogenic area in our preclinical model was caused by the ablation of satellite cells prior to ischemia, we investigated whether the increased adipogenesis in the regions of greater ischemia correlated with a loss or reduction in the number of Pax7^+^ cells. Immunostaining for Pax7 was performed on paired proximal and distal skeletal muscle sections from each subject. Although Pax7^+^ cells were still present in all subjects’ distal muscle, we observed a significant decrease in the number of Pax7^+^ cells in distal tissue (Fig. 8B). These findings support the possibility that chronic ischemia results in loss of satellite cell number and/or possibly dysfunction, which may contribute to the pathogenesis of PAD in general and CLI in particular.

**Figure 8:**
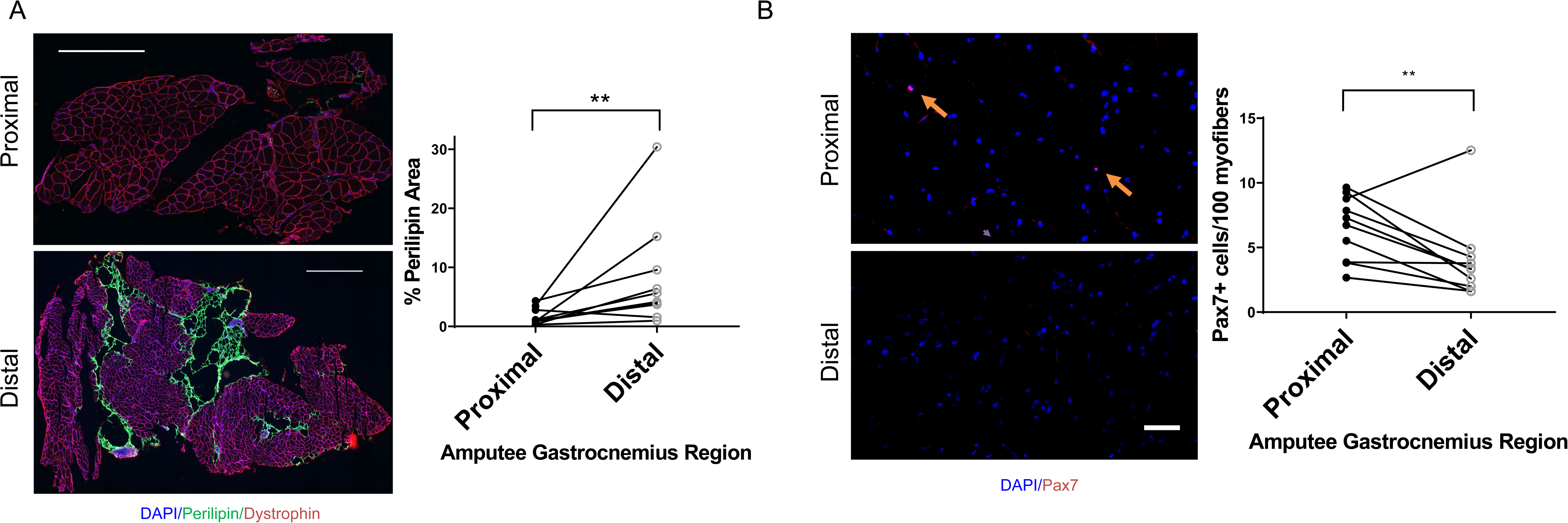
CLI patients have increased adipogenic changes in more ischemic muscle regions that correspond with decreased Pax7^+^ cell numbers. (A) Perilipin staining in the gastrocnemius muscle of CLI patients revealed significantly greater fat deposition in more distal ischemic regions. Scale bar= 1 mm (B) More ischemic distal regions of the same patients in panel (A) had significantly fewer Pax7^+^ cells, Scale bar= 100 μm

## Discussion

Although surgical and endovascular approaches to revascularization represent the primary strategy to treat PAD, outcomes remain poor, particularly in CLI, which results in high rates of subsequent amputation (30, 31). Moreover, while experimental pro-angiogenic approaches to improve limb perfusion have shown great promise in preclinical models of hind limb ischemia, they have proven suboptimal in clinical experience (32). We hypothesized that these poor outcomes might be explained, at least in part, by non-vascular etiologies of CLI. Our prior results supported this hypothesis by demonstrating that skeletal muscle cell responses to ischemia are independent of blood supply and are strongly influenced by genetic background. However, the role of skeletal muscle regeneration in the response to ischemia and, in particular, the role of muscle progenitor cells in this process, remained unknown. Here, we have demonstrated an absolute requirement for Pax7^+^ skeletal muscle satellite cells in muscle regeneration following ischemic injury. Furthermore, by continuously feeding mice a tamoxifen-containing diet over 30 days post-HLI, we ensured that there was no repopulation of the satellite cell pool (29), and we demonstrated a response to ischemia that was entirely muscle-dependent. Although one pior study raised the possibility that, following a critical juvenile period, satellite cells were dispensible for regeneration in the postnatal phase, our results are consistent with studies that demonstrate an absolute requirement for satellite cells during regeneration, in this case following ischemia-induced muscle injury (33, 34). Our data demonstrate that complete recovery from ischemia follows a similar time course as skeletal muscle injuries that are cytotoxic and cryogenic in nature (21, 23).

Staining for endothelial cells in mice lacking satellite cells verified that vascular cells were not targeted non-specifically by DTA after tamoxifen treatment and, therefore, that the observed injury was not due to loss of muscle perfusion. Somewhat surprisingly, we found that the capillary density was in fact increased in Pax7^Δ^ mice. Although the mechanisms responsible for this effect are not clear, it is possible that capillary proliferation occurred as a compensatory response to the increased tissue destruction (35). One caveat in interpreting this result is that decreased muscle area due to atrophy could have falsely increased apparent vascular density. Future studies will be necessary to fully elucidate the nature of the endothelial response during this process, including examination of endothelial cell proliferation, angiogenesis, and collateralization, which are known to occur in the setting of hindlimb ischemia (36).

Using several complementary approaches (oil red O, BODIPY, perilipin), we demonstrated the novel and important finding that in the absence of Pax7^+^ satellite cells, ischemia induces marked lipid deposition within skeletal muscle. This observation distinguishes the injury in this model from that seen in murine models of muscular dystrophy and cardiotoxin injury, which lack similar adipogenesis. Although the mdx mouse model lacks the extreme fat deposition that is observed in DMD patients (25), a “humanized” mdx model with shorterned telomeres and mitochondrial defects did show greater adipogenic changes (37). These lipid deposits are presumed to be pathogenic, because many skeletal muscle diseases result in adipogenic changes (38). Notably, the complete loss of satellite cells in Pax7^Δ^ mice recapitulated findings seen in muscle tissue samples of CLI patients, who had increased adipogenesis in more ischemic distal regions. The mice used in this study were on a C57BL/6 background, a strain in which the skeletal muscle is known to be resistant to ischemic injury (10). Strikingly, the absence of satellite cells completely abrogated the protective effect conferred by C57BL/6 genetic factors, suggesting that satellite cell loss or dysfunction contributes to the CLI phenotype. Consistent with this observation, we found that more ischemic distal regions of CLI muscle had significantly fewer Pax7^+^ satellite cells. It is important to note that the mouse phenotype was induced by the complete absence of satellite cells after tamoxifen treatment, although it is unclear whether partial loss of Pax7^+^ cells would result in a similar phenotype. Although satellite cells are present in more ischemic regions, it is possible that they may be dysfunctional and unable to contribute to regeneration. Satellite cell dysfunction may not manifest as a decrease in absolute number, but there may instead be epigenetic modifications or post-transcriptional and post-translational alterations that affect satellite cells’ ability to effectively promote regeneration in CLI patients. Alternatively, the reduction in Pax7^+^ cell number with ischemia in CLI may result from a loss due to satellite cell exhaustion reminiscent of phenotypes seen in DMD patients. Future experiments will be necessary to elucidate the exact role that satellite cells play in the pathogenesis of PAD. Gene expression profiling of satellite cells in PAD patients with claudication or CLI may identify a specific genetic signature that defines the pathophysiology of satellite cells in these conditions. The observed correlation between preclinical and clinical adipose deposition supports the biological and clinical relevance of these findings.

FAPs have been shown to play a role in obesity-associated skeletal muscle dysfunction as well as in denervated skeletal muscle (39, 40). We hypothesized that FAPs were responsible for the increased adipogenesis after ischemia in Pax7^Δ^ mice. To explore this possibility, we inhibited MMPs with batimastat, an approach that has been shown to inhibit FAP differentiation into adipocytes. Indeed, we observed a decrease in the degree of adiposity after batimastat treatment, but this was accompanied by a corresponding increase in fibrosis. Future studies, such as lineage tracing using an FAP marker such as PDGFRα (24) will be necessary to conclusively determine whether FAPs or other progenitor cell types contribute to this fat infiltration.

Several important questions arise regarding the mechanisms responsible for both the adipogenesis and the switch to a fibrotic phenotype after batimastat treatment. First, what are the paracrine signaling pathways between satellite cells and other muscle progenitor cells, including FAPs, that drive normal myogenesis? Pax7^+^ cells account for a small percentage of total cells in muscle tissue, yet in typical muscle cell isolates, the vast majority of mononuclear cells have the capacity to fuse and differentiate into myotubes *in vitro*, suggesting that the presence of satellite cells confers on other MPCs (e.g., myoblasts, pericytes, FAPs) the ability to differentiate into functional muscle. This likely involves paracrine signaling mechanisms that remain to be fully elucidated, although PDGF-BB and DLL4 have been implicated in driving pericytes toward a myogenic lineage (41). Second, what are the mechanisms that drive the increased adipogenesis in the absence of satellite cells? Does a suppressive signal from satellite cells to FAPs normally prevent adipogenesis, or does the absence of satellite cells activate another pathway to drive adipogenesis? Third, and equally important, does ischemia contribute to these processes, since adipogenesis does not occur in the non-ischemic limb, or are these pathways driven by aberrant regeneration? Future studies will be necessary to elucidate these mechanisms, and it is hoped that such information would lead to the eventual development of therapies for diseases of aberrant muscle stem cell number and/or function, such as CLI and DMD. Batimastat provides a potential starting point for development of drugs to inhibit adipogenic changes in skeletal muscle. Although an increase in fibrosis in CLI in place of adipose tissue may not translate into better clinical outcomes, it provides an initial strategy to prevent pathological adipogenesis.

## Methods

### Mouse lines and tamoxifen treatment

For satellite cell genetic ablation experiments, Pax7-Cre^ERT2^ mice (Jackson Labs Stock: 017763, B6.Cg-*Pax7^tm1(cre/ERT2)Gaka^*/J) were backcrossed to ROSA26^DTA^ mice (Jackson Stock: 009669, C.129P2(B6)-*Gt(ROSA)26Sor^tm1(DTA)Lky^*/J). Sterile-filtered tamoxifen (Sigma) or corn oil at 75 mg/kg body weight was given to each mouse via an intraperitoneal route for 5 days. Following five days of treatment with either tamoxifen or control, mice were given tamoxifen or control matched diet (Teklad Tamoxifen Diet from Envigo and 2016 Control Diet from Teklad) to continue treatment at a lower dose of 50 mg/kg body weight. All mice were used at 8-12 weeks unless stated otherwise. All protocols described here have been approved by the Duke University Institutional Animal Care and Use Committee and are consistent with the guidelines published by the United States Department of Health and Human Services.

### Hindlimb Ischemia Surgery and Perfusion Imaging

Hindlimb Ischemia surgery was performed as previously described by our lab.(10, 42) Briefly, mice were anaesthetized on a heated pad using 1.5 L/minute oxygen and 1-3% isoflurane. Prior to surgery, the mice were scanned with a Laser Doppler Perfusion Imager (LDPI from Moor Instruments) to obtain a baseline perfusion in the hindlimbs. Using sterile surgical instruments (sterilized by autoclaving), a 1 cm incision was made just below the inguinal ligament. Subcutaneous fat was then cleared and the femoral artery was separated from the neurovascular bundle carefully without perforating the femoral vein. A 7-0 silk non absorbable suture (Sharpoint) was used to ligate the femoral artery above the bifurcation of the lateral circumflex femoral artery and a ligature was also made below superficial caudal epigastric artery but above the bifurcation of the popliteal artery. The wound was then closed using an absorbable Vicryl 5-0 suture (Ethicon). A post-surgery LDPI perfusion scan was then performed on the animal to verify complete closer of the artery. The animals were provided appropriate pain relief and monitored after surgery to ensure animal welfare.

### Tissue Collection and Muscle Processing

Mice were euthanized using approved euthanasia protocols, and the tibialis anterior (TA) and extensor digitorum longus (EDL) muscles were isolated and frozen using liquid nitrogen in Optimal Cutting Temperature (OCT) medium, while the gastrocnemius muscle was flash frozen for tissue analysis at −80 degrees Celsius. 8 micron sections of tissue were obtained by sectioning on a Leica 3150S cryostat at −25 degrees Celsius and stored on Superfrost Slides.

### Immunofluorescence, Microscopy and Image Analysis

Following sectioning and storage of slides, slides were retrieved from −80 degrees Celsius and allowed to equilibrate to room temperature. Slides were then fixed in 4% paraformaldehyde, followed by permeabilization in 0.2% Triton X in Phosphate Buffered Saline (PBS). After washes in PBS, the slide was blocked with 5% goat serum in PBS for 1 hour. For satellite cell staining, a mouse IgG blocking antibody (Jackson Immunoresearch) was used in the blocking buffer and an antigen retrieval step was performed by heating the slides in a pressure cooker in citrate buffer was also performed prior to incubation with the primary antibody. The antibody of interest was then incubated at the dilutions listed in the table below overnight at 4 degrees Celsius. The following day, slides were washes 3 times in PBS, followed by incubation of the species matched secondary antibody (Alexa Flour conjugated) and incubated at room temperature for 1 hour. The slide was then either washes 3 times with PBS or PBST, and if needed, a 5 minute incubation of a nuclear stain (either DAPI or Hoechst stain) was added and then washed. Slides were then mounted in either Vectashield mounting medium or Prolong Gold mounting medium and allowed to cure overnight. Brightfield microscopy and epiflourescent microscopy were both performed on a Zeiss Upright AxioImager, while all confocal microscopy was performed on the Zeiss 780 or Zeiss 880 Inverted Confocal Microscope. All image analysis were performed in Zeiss Zen software, IMARIS or ImageJ with identical thresholds and blinding performed for all signal quantification.

### Hematoxylin and Eosin Staining

Sections were brought to room temperature, then fixed for 10 minutes in 10% Neutral Buffered Formalin (NBF). The slides were then washed with distilled water, followed by staining of nuclei with the Meyer’s haematoxylin for 4 minutes. Slides were then rinse in running tap water for 10 minutes and then differentiated with 0.3% acid alcohol. An additional rinse in tap water and in Scott’s tap water substitute was used to further enhance coloration of the nuclei. Samples were then briefly incubated in 90% Ethanol (EtOH) and then stained with alcoholic eosin for 30 seconds. The slides were then dehydrated in 100% EtOH followed by 2 rinses in Xylene or Xylene substitute for 2 minutes each before mounting with Cytoseal resin based mounting medium. The slides were then allowed to cure overnight before imaging.

### Muscle Contractile Experiments

Contractile muscle force was measured as described previously(43). Briefly, single EDL muscles were isolated and ligated with a 5–0 silk suture at each tendon and maintained in a physiological saline solution (pH 7.6) containing (mM) 119 NaCl, 5 KCl, 1 MgSO4, 5 NaHCO3, 1.25 CaCl2, 1 KH2PO4, 10 Hepes, and 10 dextrose at 30□C under aeration with 95% O2–5% CO2 throughout the experiment. The muscle was mounted in a bath within the force transducer (Aurora 300B-LR) operated in isometric mode. A five minute equilibration step was performed, during which single twitches were elicited every 30 s with 0.5 ms electrical pulses. Isometric tension was evaluated by 250 ms trains of pulses delivered at 10, 20, 40, 60, 80, 100 and 120 Hz. After the experimental protocol, muscle length was determined with a digital caliper and muscle mass was measured after removing liquid. The cross-sectional area for each muscle was determined (mass of the muscle (g) divided by the product of its length (Lo, mm) and the density of muscle and was expressed as square millimetres. Muscle output was then expressed as isometric tension (N cm−2) determined by dividing the tension (N) by the muscle cross-sectional area. In the case of atrophied muscle, absolute tension was used as the measure of force because the cross sectional muscle area is no longer an accurate measure as the density is now changed.

### Oil Red O Staining

Frozen tissue sections were first air dried to the slides at room temperature. After fixing in formalin for 4 minutes, the samples were briefly washed under running tap water for 1 to 10 minutes. After a rinse with 60% isopropanol, samples were stained with freshly prepared Oil Red O working solution (Oil Red O Powder from Sigma in 60% isopropanol) for 15 minutes, then rinsed again with 60% isopropanol. The samples were then lightly stained with Meyer’s haematoxylin and rinsed with distilled water. Slides were mounted in aqueous glycerine jelly and imaged within 2 hours.

### BODIPY staining

Frozen sections were fixed with 4% PFA in PBS for 10 minutes, followed by 2 washes with PBST for 5 minutes each. The slides were then incubated for 60 min with 1 ug/mL BODIPY 493/503 in PBST. Following incubation, the slides were washed twice with PBST for 5 minutes each, then twice with PBS for 5 minutes each. Prolong Diamond with DAPI was applied and the slide was imaged immediately.

### microCT/DiceCT

EDLs were isolated from hindlimb of mice and fixed immediately in 10% NBF solution overnight. The muscles were then stained using the diceCT protocol as described previously (44). Briefly, the muscles were incubated in Lugol’s Iodine for 2 nights. The muscles were then scanned in a fixed container at low power in a Nikon XTH 225 ST microCT scanner at a 10 micron or lower resolution. Images were then reconstructed on the proprietary Nikon automated reconstruction software and analyzed using Avizo to delineate soft tissue densities in false colors.

### Batimastat Treatment

Pax7-Cre^ERT2^; ROSA26^DTA^ mice were all given tamoxifen injections at their usual dose of 75 mg/kg body weight for 5 days prior to surgery. 1 day prior to surgery, half the mice were given the drug Batimastat at 30 mg/kg body weight as a suspension of 3 mg/ml in sterile filtered phosphate-buffered saline containing 0.01% Tween 80 and the other half was given a vehicle control without the Batimastat. The injections were then given daily until muscle was isolated for freezing and sectioning. Following surgery, the mice were switched to a tamoxifen diet and the muscle was harvested 7 days post surgery.

### Fast Green/Sirius Red Staining

Samples were first placed in 0.04% Fast Green (Sigma) for 15 min, then washed with distilled water. They were then incubated in 0.1% Fast Green and 0.04% Sirius Red (Sigma) in saturated picric acid for 30 min. Then, samples were dehydrated through 70%, 90% and 100% Ethanol and cleared in Xylene for 2 minutes before mounting using Cytoseal mounting medium. Positive and negative controls were ran to validate the specificity of this assay for collagen.

### Myoblast isolation

Hindlimb muscles from mice were dissected and rinsed briefly in sterile PBS, and placed into p100 dish containing DMEM + 1% p/s. From this point on, all steps were performed in a Biological Safety Level 2 hood under sterile conditions. Muscles were cleaned of excess connective tissue and tendons and transferred to a 2nd p100 dish containing 5 mL DMEM + 1% p/s. The muscle was then minced into fine pieces using razor blades held with hemostats for over 10 min, then transferred to a 50 ml centrifuge tube using a wide-bore pipet. The pellet was then centrifuged using a tabletop centrifuge for 2 minutes at speed 800g. The medium was aspirated and 18mL DMEM + p/s was added to loosen pellet. 2ml of 1% pronase to bring the final volume to 20ml was added (0.1% pronase final) and the mixture was digested at 37 C for 1 hour in a 50 mL conical on the Nutator. The cells were then spun down for 3 min at 800g and medium aspirated. The muscle was suspended in 10 mL DMEM + 10% FBS + p/s and triturated many times (20) to loosen cells. The supernatant was put through a Steriflip 100μm vacuum filter and the filter was washed with 5ml of DMEM + 10% FBS + p/s. The cells were then spun for 5 minutes at speed 1000g and resuspended in 10 mL of growth media (Ham’s F10 w 20% FBS and P/S) and plated on collagen coated plates.

### In vitro Myogenic and Adipogenic Differentiation

Differentiation of isolated myoblasts was stimulated by plating the cells on entactin-collagen-laminin coated plates in conditions of serum withdrawal in differentiation medium (DMEM supplemented with 2% horse serum, 1% penicillin/streptomycin, 0.2% amphotericin B, and 0.01% human insulin/transferrin/selenium). The cells were placed either in normoxic conditions in the incubator or in a hypoxic (0% oxygen) chamber. Media was changed daily to ensure cell viability.

Adipogenic differentiation was induced by placing cells for 48 hours in medium containing 10% FBS, 0.5 mM isobuylmethylxanthine, 125 nM indomethacin, 1 μM dexamethosone, 850 nM insulin, 1 nM T3 with or without 1 μM rosiglitazone (Sigma). After 48 hours, cells were switched to medium containing 10% FBS, 850 nM insulin, 1 nM T3 with 1 μM rosiglitazone. Cells were placed in either normoxic or hypoxic conditions as described above, and media was changed every other day to ensure cell viability.

### Human Skeletal Muscle Biopsy

Critical Limb Ischemia (CLI) patients undergoing above-knee or below knee amputations were consented under an approved Institutional Review Board (IRB) protocol to donate skeletal muscle tissue from the amputated limb. Muscle biopsies were collected from both the proximal and distal ends of the gastrocnemius muscle and oriented in a cross sectional manner in OCT and frozen in liquid nitrogen. The samples were then sectioned in a cryostat and stored at −80 C for subsequent analysis.

### Statistical Analysis

For each of the analysis, a script was used to blind the reviewer to the either the images or the murine treatments to ensure that there is no bias in analysis. For a comparison of 2 groups, a two-way student’s t-test was performed in GraphPad Prism and statistical significance was established at p<0.05. For multiple group comparisons, an ANOVA was first performed to determine whether an effect was present, following by a t-test for multiple groups with a correction for multiple group testing in GraphPad Prism. Significance was once again established by a corrected p value <0.05.

**Table 1:** Commercially available antibodies used for immunofluorescence.

| **<u>Antibody</u>** | **<u>Manufacturer</u>** | **<u>Catalog #</u>** | **<u>Dilution</u>** | **<u>Species</u>** |
| --- | --- | --- | --- | --- |
| Pax7 | DSHB | Pax7 | 1:50 | Mouse IgG1 |
| Dystrophin | Thermo | RB-9024 | 1:100 | Rabbit |
| Type IIa<br>Fibers | DSHB | SC-71 | 1:100 | Mouse IgG1 |
| Type I<br>Fibers | DSHB | BA-F8 | 1:100 | Mouse<br>IgG2b |
| Type IIb<br>Fibers | DSHB | BF-F3 | 1:100 | Mouse IgM |
| Embryonic<br>Myosin<br>Heavy Chain | DSHB | F1.652 | 1:50 | Mouse IgG1 |
| CD31 | BioRad | MCA2388 | 1:200 | Rat |
| Dystrophin | Abcam | ab3149<br>(discontinued) | 1:100 | Mouse IgG1 |
| Perilipin | Cell Signaling<br>Technologies | 9349S | 1:200 | Rabbit |
| Myogenin | DSHB | F5D | 1:50 | Mouse IgG1 |
| Myosin<br>Heavy Chain | DSHB | A4.1025 | 1:200 | Mouse<br>IgG2a |

## Author contributions

HA and CDK designed the research study. HA conducted all *in vivo* and *in vitro* experiments and performed data analysis. LAO, BFG, and KWS isolated human skeletal muscle and assisted in human muscle staining and experiments. MP performed animal husbandry, genotyping and HLI surgeries. CES and JM conducted muscle force generation experiments. HA wrote the manuscript, and CDK co-wrote and edited the manuscript.

## Acknowledgments

We would like to thank the Light Microscopy Core Facility at Duke University for the use of their microscopes and assistance in image acquisition; the Shared Materials Instrumentation Facility for the use of the microCT scanner and assistance in data acquisition and analysis; Drs. Mitchell Cox, Cynthia Shortell, Chandler Long and the Division of Vascular Surgery in the Department of Surgery for their assistance in acquiring human skeletal muscle samples; Timothy J. McCord, for his expert consultation in imaging and image analysis; and Jianbin Li for assistance with animal husbandry. This study was supported in part by NIH grants HL124444 and HL118661 (to CDK) and HL125695 (to JMM).

---

**Figure S1.**
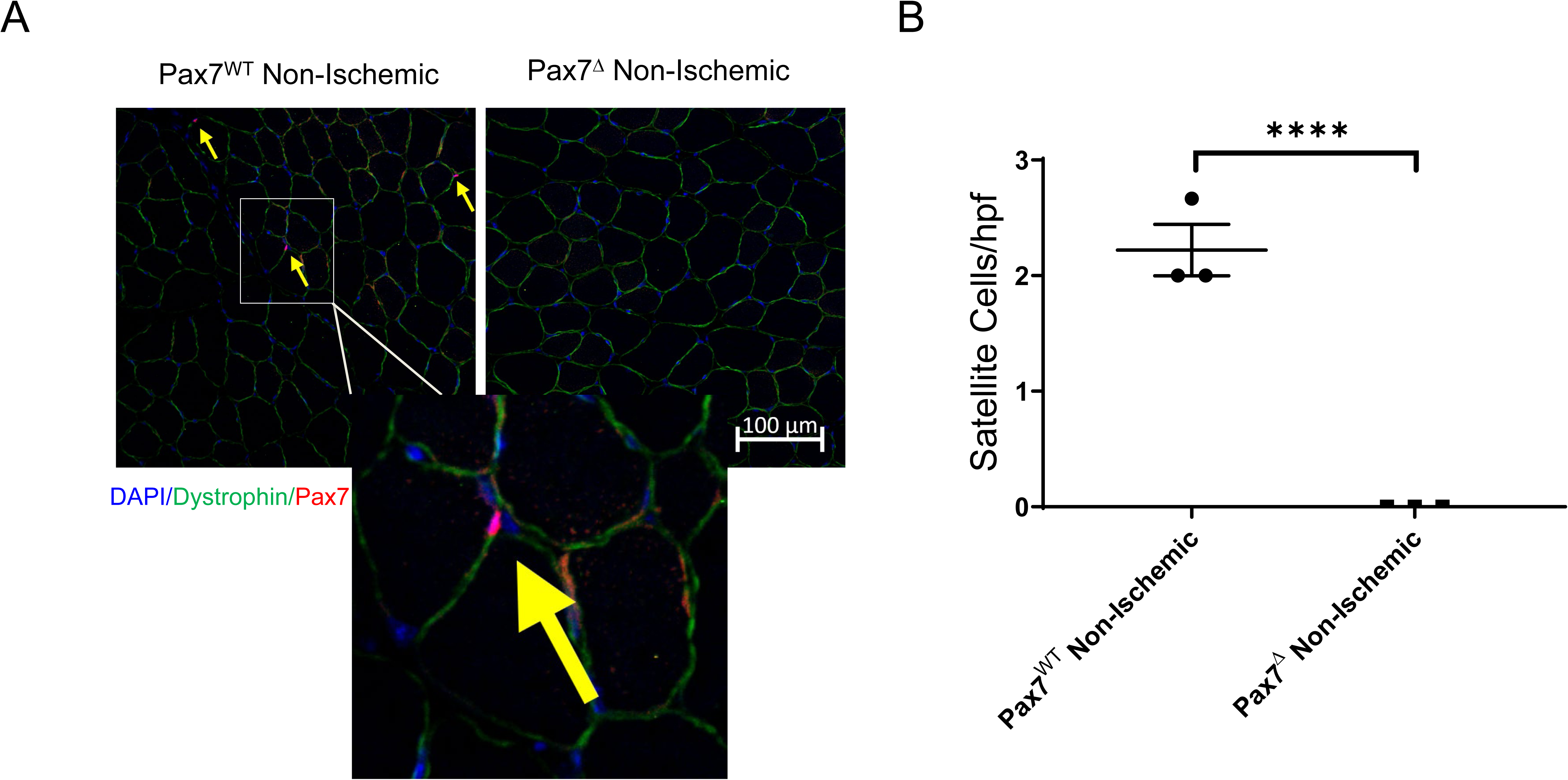
Pax7^CreERT2^;ROSA26^DTA^ mice display a complete loss of satellite cells in resting muscle after tamoxifen treatment. (A) Pax7 immunostaining of the non-ischemic TA muscle revealed typical satellite cells at the periphery of resting skeletal muscle, directly beneath the basal lamina. No Pax7 staining was observed in tamoxifen-treated mice. (B) Quantification of satellite cell numbers demonstrated a complete absence of satellite cells following tamoxifen treatment. Scale bar=100 μm

**Figure S2:**
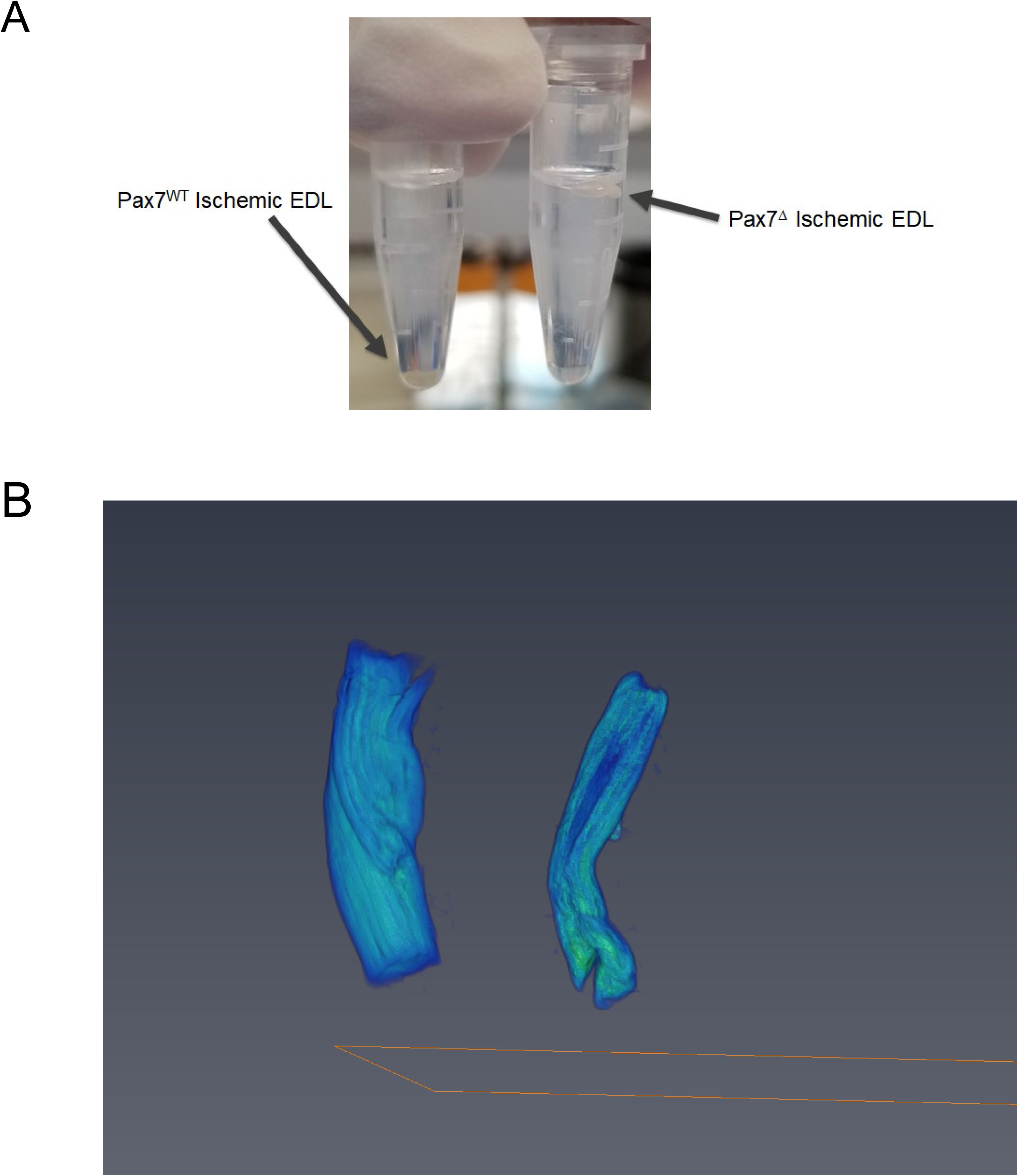
Tissue density is reduced in Pax7^Δ^ EDLmuscle 30 days after HLI due to adipogenic changes. (A) Thirty days after HLI, Pax7^WT^ ischemic EDL muscle sinks in aqueous fixative, while the ischemic Pax7^Δ^ muscle floats, suggesting a large difference in soft tissue density due to fat accumulation in the Pax7^Δ^ muscle. (B) diceCT of 30-day-ischemic Pax7^WT^ EDL muscle (left) showed individual, regenerated, fused muscle fibers while ischemic Pax7^Δ^ muscle had significant soft tissue fatty changes 30 days after ischemia as well as general muscle atrophy, visualized in green. (C) Video from diceCT of EDL muscle 14 days after ischemia showed significant soft tissue fatty changes, shown in green, in Pax7^Δ^ ischemic muscle.

